# Exact Probability Distribution for the ROC Area Under Curve

**DOI:** 10.1101/2022.12.13.520256

**Authors:** Joakim Ekström, Jim Ögren, Tobias Sjöblom

## Abstract

The Receiver Operating Characteristic (ROC) is a *de facto* standard for determining the accuracy of *in vitro* diagnostic (IVD) medical devices, and thus exactness in its probability distribution is crucial toward accurate statistical inference. We show the exact probability distribution of the ROC AUC-value, hence exact critical values and p-values are readily obtained. Because the exact calculations are computationally intense, we demonstrate a method of geometric interpolation which is exact in a special case but generally an approximation, vastly increasing computational speeds. The method is illustrated through open access data, demonstrating superiority of 26 composite biomarkers relative to a predicate device. Especially under correction for testing of multiple hypotheses, traditional asymptotic approximations are encumbered by considerable imprecision, adversely affecting IVD device development. The ability to obtain exact p-values will allow more efficient IVD device development.

## 1 BACKGROUND

The ROC concept^1^ has become a *de facto* standard for determining the accuracy of binary predictors within many areas, such as IVD medical devices^2,3,4^. In order to provide a regulator with valid scientific evidence supporting the conclusion that there is reasonable assurance that an IVD device is safe and effective^5^, it is highly valuable to demonstrate that the ROC Area Under Curve (AUC) value of the novel IVD device is non-inferior relative to a predicate device. Because the ROC-curve and the AUC-value is subject to randomness due to sources of error including sampling error and measurement imprecision, statistical hypothesis tests are necessary.

The motivation for this work, specifically, is our effort to develop biomarkers for early detection of cancer. Because of combinatory proliferation when composite biomarkers are generated, the correction for testing of multiple hypotheses is often severe; yielding critical values that are situated in the far tails of the AUC-value probability distribution. It is a well-known phenomenon that the tails of probability distributions are particularly sensitive to inaccuracies; hence exactness in the AUC-value probability distribution is crucial toward accurate statistical inference.

The term ROC was introduced by^6^ and the phrase AUC was used by^7^; Chapter 1 of^8^ contains historical notes. Work on the AUC-value distribution has been conducted through an assumption of asymptotic normality, allowing a t-test or similar. In particular,^9,10,11^ make inference through, respectively, the Wilcoxon statistic, the Wilcoxon statistic with correlation, and a binormal distribution. Articles^12,13^ make inference through the F-test and a binormal distribution. The work^14^ proposes a new AUC variance estimate and applies it toward sample size estimation, and^15^ argues in favor of Monte Carlo-simulation of AUC-values. Reviews^16,17^ contain historical notes.

## 2 PROBABILITY DISTRIBUTION OF THE AUC-VALUE

### 2.1 Exact computation via order statistics

This subsection derives the exact AUC-value probability distribution under the assumption of observed values of the True Positives (TPs) and True Negatives (TNs) that each are independent and identically distributed, *iid* with probability distribution functions denoted *F* and *G*, respectively.

Because the ROC-curve is determined by the ranks of the observed values of the TPs and TNs, the probability of an AUC-value can be computed through order statistics. In general, suppose there are *n* observed values of the TPs, *x*_1_, …, *x*_*n*_, and *m* observed values of the TNs, *y*_1_, …, *y*_*m*_, and the two have probability density functions *f* and *g* respectively, then under the *iid* assumption the joint probability density function of the order statistic, *η*, presuming they are indexed such that *y*_1_ ≤ … ≤ *y*_*m*_ and *x*_1_ ≤ … ≤ *x*_*n*_, equals

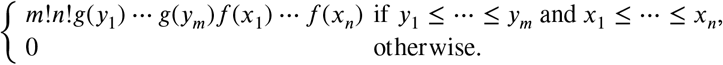

Because each ROC-curve is determined completely by the ranks of the observed values of the TPs and TNs, the probability that a given ROC-curve will manifest itself can thus be obtained through integration of the joint probability density function.

For example, the ROC-curve that has AUC-value equal to unity corresponds to observing values of the TPs and TNs such that

*y*_*m*_ ≤ *x*_1_, and under the *iid* assumption and existence of probability density functions, the probability of the ROC-curve equals

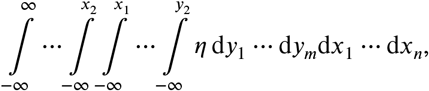

which after application of the chain rule can be shortened to

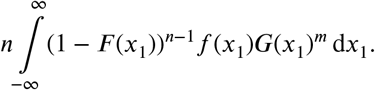

For another example, the ROC-curve that has AUC-value equal to 1 − 1/*mn*, i.e. the second highest AUC-value possible, corresponds to observing values of the TPs and TNs such that *y*_*m*−1_ ≤ *x*_1_ ≤ *y*_*m*_ ≤ *x*_2_, and under the same assumptions the probability of that ROC-curve equals

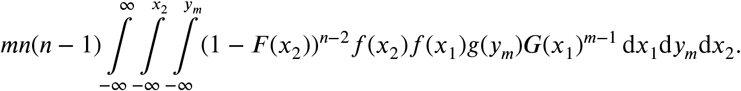

Continuing to the example of the ROC-curve with AUC-value 1 − 2/*mn*, the reader will note that there are two distinct ROC-curves that produce the AUC-value, one that corresponds to observing values of TPs and TNs such that *y*_*m*−1_ ≤ *x*_1_ ≤ *x*_2_ ≤ *y*_*m*_ ≤ *x*_3_ and the other such that *y*_*m*−2_ ≤ *x*_1_ ≤ *y*_*m*−1_ ≤ *y*_*m*_ ≤ *x*_2_. Because the two ROC-curves are mutually exclusive, the probability of observing TPs and TNs such that the AUC-value equals 1 − 2/*mn* is equal to the sum of the probabilities of those two ROC-curves. Generally, distinct ROC-curves are mutually exclusive and consequently the probability of a given AUC-value equals the sum of the probabilities of all ROC-curves that produce the given AUC-value.

In practice, numerical computation requires explicit probability distribution functions *F* and *G*. Because the ROC-curve is determined by the ranks only, it is the experience of the authors that many choices of symmetric distributions *F* and *G* of equal inter-quartile range tend to yield relatively similarly shaped ROC-curves. In any case, one example of choices of probability distribution functions are two normal distributions with equal variance and some difference in their means. Since many common parametric choices of probability distributions are quite smooth, numerical integration using the trapezoid method will often be quite accurate and also fast in its algorithmic implementation^18^.

The probability distribution function of the AUC-value can be determined by computing the probability for each of the AUC-values 0, 1/*mn*, 2/*mn*, …, 1; for an illustration see Figure 2a. However, the number of ROC-curves that produce a given AUC-value tends to be large in many situations, particularly when the AUC-value is around the center of the unit interval, and consequently determining the whole AUC-value probability distribution function through this method is in many instances impractical computationally.

### 2.2 Monte Carlo-simulation of AUC-value probabilities

When estimated through *iid* sampling, by the strong law of large numbers the empirical distribution function converges point-wise, with probability one, to the distribution function from which the observations were drawn. The result can be strengthened further; through the Glivenko-Cantelli lemma the convergence is uniform and through Donsker’s theorem the normalized difference converges in distribution to a Gaussian process^19^. Because of its desirable asymptotic properties, and properties such as simplicity of computation, the empirical distribution function is commonly utilized for estimation of distribution functions through Monte Carlo-simulation.

Percentiles can be estimated through the empirical distribution function by taking the infimum of the superlevel set, i.e. if *α* is a number in the unit interval then the 100*α*-percentile is estimated through inf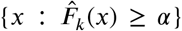, where 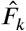 denotes the empirical distribution function at sample size *k*. In a common algorithmic implementation, determining the empirical distribution function and the infimum of the superlevel set amounts to sorting the observed values and selecting the value that is the 100*α* percent largest. Through the aforementioned beneficial asymptotic properties of the empirical distribution function, the percentile estimate obtains many desirable properties, however if the distribution function is constant in an interval then the percentile estimate obtains a discontinuity when viewed as a function of *α*. Further, even if the distribution function is strictly increasing, a relatively shallow slope of the distribution function yields the practical problem of a slow rate of convergence.

When correction for testing of multiple hypotheses is applied, which is common in for instance biomarker discovery, the critical values to be estimated are typically situated in the far tail of the probability distribution, where the distribution function often has a shallow slope. As an illustration, if the type-one error probability subsequent to correction for testing of multiple hypotheses equals 10^−10^, then the Monte Carlo-simulation estimate of the corresponding critical value is equal to the maximum AUC-value simulated for all simulations that are constituted of 10^10^ or fewer simulated AUC-values. But on contrast, as discussed in Section 2.1, computing the exact probability of AUC-values is relatively quick when the AUC-values are close to the extremities of the unit interval. A pragmatic compromise is to employ exact computation in the far tails and use Monte Carlo-simulation in the body of the AUC-value distribution.

### 2.3 Geometric interpolation

As discussed, the number of AUC-value probabilities that can be computed exactly is in practice limited by availability of computational resources. In the experience of the authors, computation of about 50 to 100 consecutive AUC-value probabilities is typically feasible when starting from zero or one. Ideally, the sum of those AUC-value probabilities is such that the difference relative to unity constitutes a number for which the corresponding percentile is suitable for estimation through Monte Carlo-simulation. However, in many instances the sum of the computed AUC-value probabilities is materially smaller than would be desired vis-à-vis percentile estimation through Monte Carlo-simulation. Consequently, the limitation of computational resources effectively yields a gap between the sum of the computed AUC-value probabilities and the largest value deemed suitable for percentile estimation through Monte Carlo-simulation. In this instance, the authors have observed a perceived stability of ratios of subsequent differences of AUC-value probabilities, arising from a special case, and exploited it toward bridging the aforementioned gap through geometric interpolation.

Denote by *i* = 0, 1, 2, 3, … the sequence of AUC-values 1, 1 − 1/*nm*, 1 − 2/*nm*, 1 − 3/*nm*, …, where *n* and *m* are, as in Section 2.1, the number of observed values of TPs and TNs respectively. Suppose the observed values of TPs are *iid* with probability distribution function *F* and the observed values of TNs are *iid* with probability distribution function *G*, and denote by *x*_0_, *x*_1_, *x*_2_, *x*_3_, … the probabilities that the AUC-value attains 1, 1 − 1/*nm*, 1 − 2/*nm*, 1 − 3/*nm*, ….

Consider firstly the special case when *F* = *G*, i.e. the observed values of the TPs and TNs follow the same probability distribution. Then the expressions of Section 2.1 simplify so that the probability of each ROC-curve equals *n*!*m*!/(*n* + *m*)!, and therefore the probability of an AUC-value is determined by the number of distinct ROC-curves that yield the AUC-value. Consequently, in this special case it holds that *x*_*i*_ = *k*_*i*_*n*!*m*!/(*n* + *m*)!, where *k*_0_, *k*_1_, *k*_2_, *k*_3_, … denotes the number of distinct ROC-curves that yield the AUC-values 1, 1 − 1/*nm*, 1 − 2/*nm*, 1 − 3/*nm*, …. Figure 1 illustrates the number of ROC-curves that yield the AUC-value 26/30 when *n* = 5 and *m* = 6; in the example *k*_4_ = 5.

**Figure 1.**
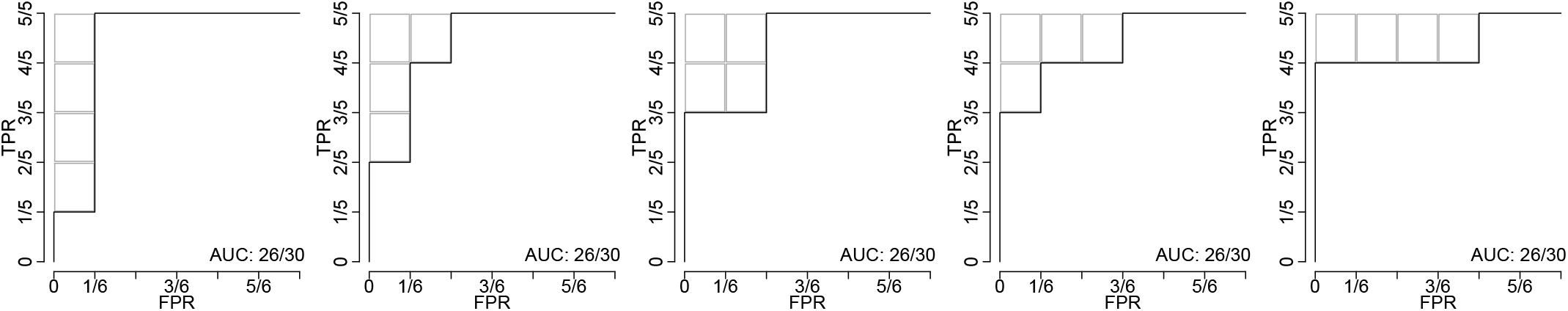
Five ROC-curves yielding the same AUC-value. Illustration of the 5 distinct ROC-curves that yield the AUC-value 26/30 when the number of TPs and TNs are *n* = 5 and *m* = 6, respectively. The gray boxes illustrate the subtracted four rectangles each with area 1/*nm*; hence 1 − 4/*nm* = 26/30.

Let Δ denote the difference operator, Δ*x*_*i*_ = *x*_*i*_ − *x*_*i*−1_. The present authors have observed that, in the top percentile of the probability distribution, the ratio of subsequent differences Δ*x*_*i*_/Δ*x*_*i*−1_ tend to be relatively well approximated by *C*Δ*k*_*i*_/Δ*k*_*i*−1_ where *C* is a constant. As discussed, in the special case when *F* = *G* equality holds with *C* = 1. See Figures 2c and 2d for a numerical example with a selection of values of *n* and *m*, and where *F* and *G* are each normal distributions with equal variance and a selection of differences in their means. Rearranging terms yields the approximation

**Figure 2.**
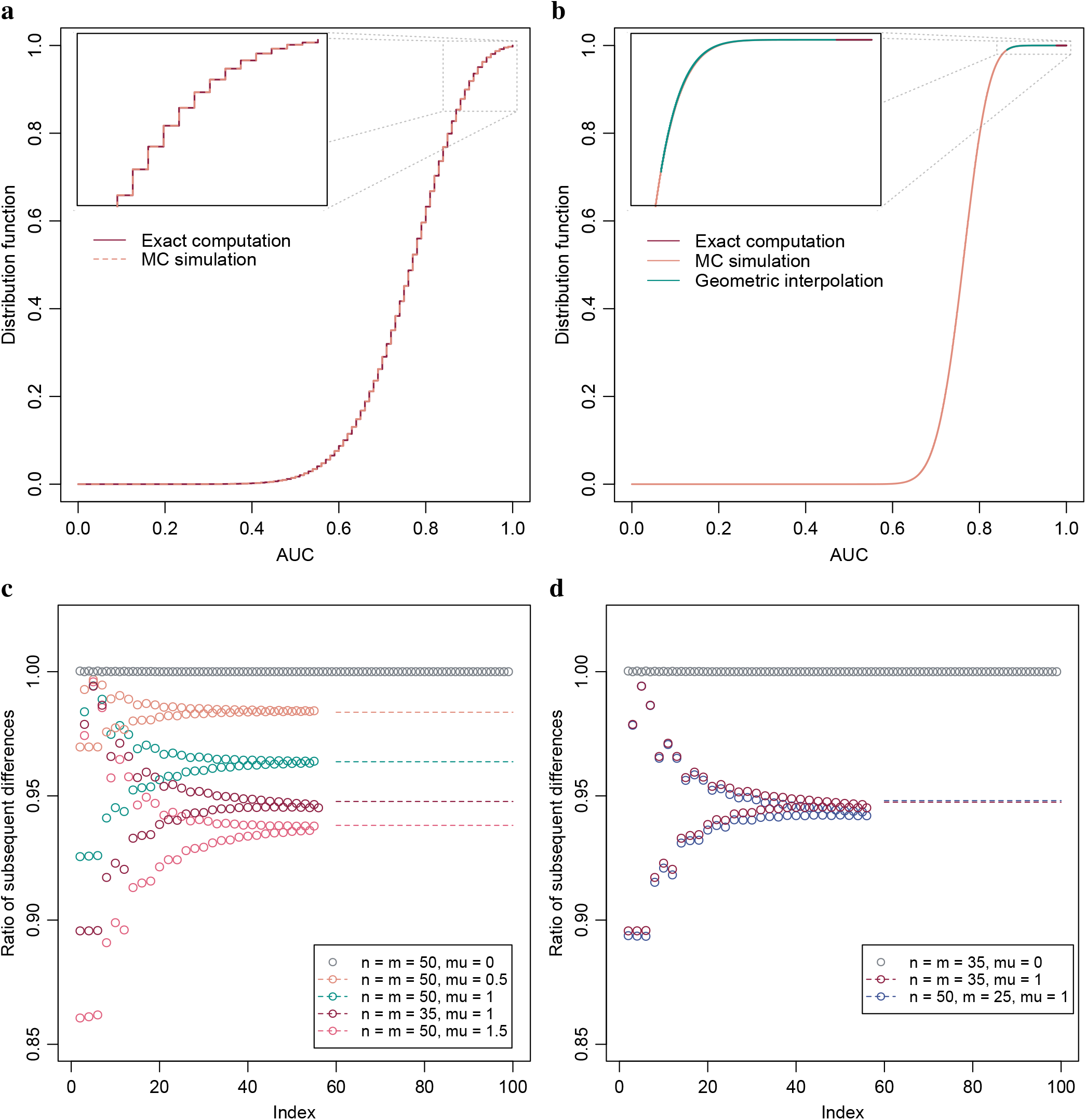
Exact probability distributions of the AUC-value under a range of parameters. (a) Exact computation and Monte Carlo estimation of the AUC-value distribution when TPs and TNs are 10 each and *iid* with normal distributions of equal variance and unit difference in mean. Inset, top 10 percentiles. (b) Exact computation and MC estimation of the AUC-value distribution when TPs and TNs are 50 each and *iid* normal with equal variance and unit difference in means. Geometric interpolation between exact computation and the MC estimated 99^th^ percentile is included. Inset, top percentile. (c) Stability of the ratio of subsequent differences under a range of parameters, i.e. (Δ*x*_*i*_/Δ*x*_*i*−1_)/(Δ*k*_*i*_/Δ*k*_*i*−1_), with solution through bisection method included in dashed lines. (d) Plot similar to Subfigure c, albeit with numbers of TPs and TNs selected such that their products are approximately equal, with two selections having equal numbers of TPs and TNs and the third twice the TPs relative to TNs.

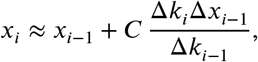

which can be employed to bridge the gap between the exact computations discussed in Section 2.1 and the Monte Carlo simulations discussed in Section 2.2.

An estimate of *C* in the above approximation can be obtained by solving for equality between the Monte Carlo estimate of the, say, 99.9^th^ percentile and the same percentile when estimated through exact computation and geometric interpolation using the above approximation. For example, the bisection method is a simple yet effective method to solve for *C*, thus obtaining an estimate using the aforementioned equality^18^. The reader will note that there will be division by zero in the above approximation when Δ*k*_*i*−1_ = 0, i.e. *k*_*i*−1_ = *k*_*i*−2_, however the authors have only seen this occur when *i* = 2 or in some instances when *i* ≈ *nm*/2 and those instances do not constitute the typical intervals of application of the approximation, cf. Figure 2b.

While analyzing the data set of^20^, see Figure 3, we noted that the aforementioned ratio is less stable when the numbers of TPs and TNs are highly unbalanced. In Figure 3d, where the balance of TPs to TNs for the pancreatic cohort is approximately 1 : 9, a slightly positive trend can be visually perceived, and when a constant is assumed the distribution function will, seen from the right, firstly drop too quickly and then too slowly, see Figure 3b. When an approximation is unsatisfactory, common approaches are to either use a more sophisticated approximation or to lessen the use of the approximation. Figure 3d shows an approximation using a monotonically increasing function, an exponent function, that connects the AUC-value probabilities with the AUC-value probability at the median which in inferred from the curvature of the S-shape yielding Δ*x*_*i*_/Δ*x*_*i*−1_ = 1. Lessening the use of the approximation is achieved by shortening the gap to be bridged; computing more exact AUC-value probabilities and estimating a higher percentile through Monte Carlo-simulation.

**Figure 3.**
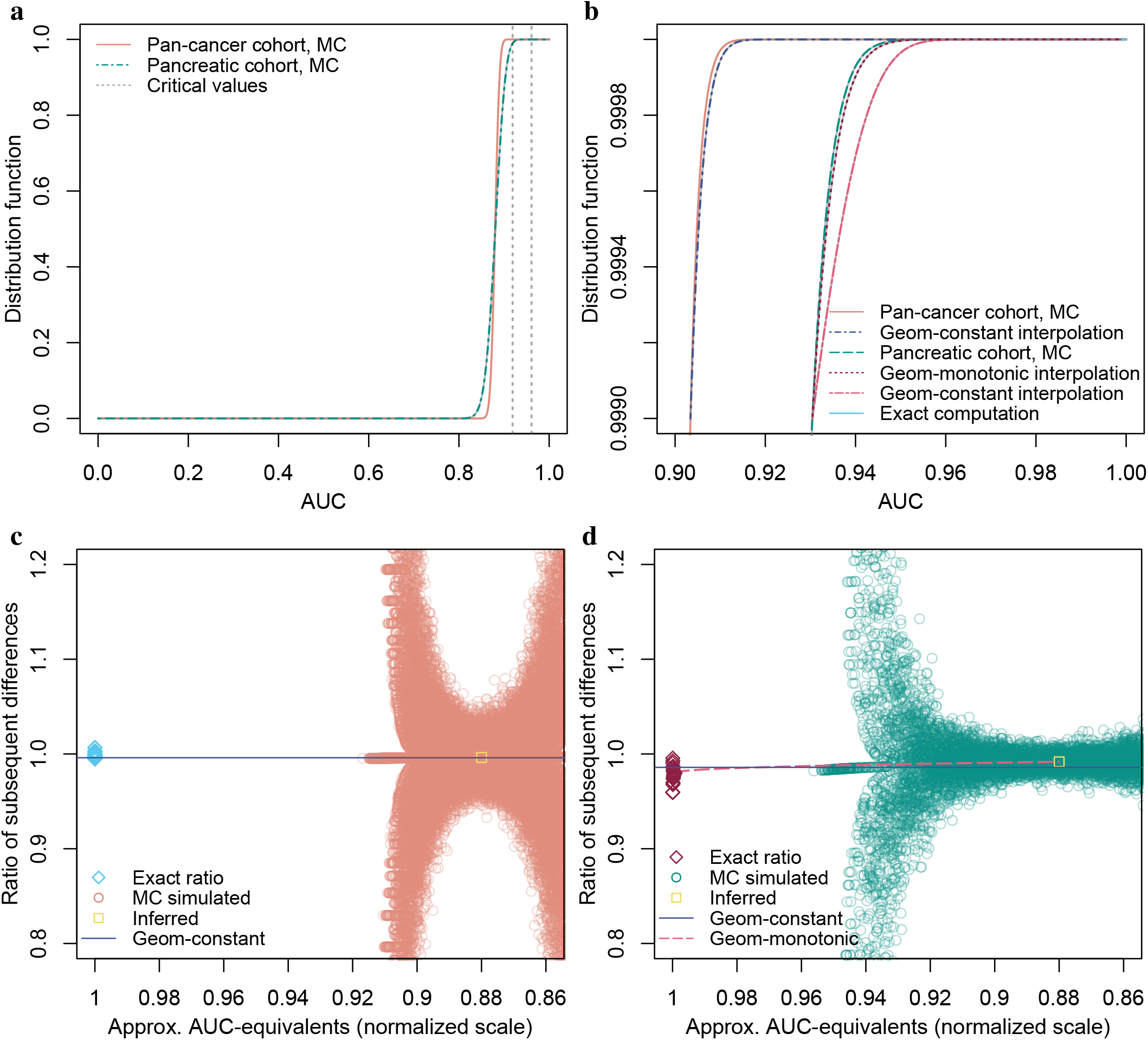
Geometric interpolation for cohorts with balanced and highly unbalanced numbers of TPs and TNs. (a) Monte Carlo-simulated AUC-value distribution functions for biomarkers on par with benchmark predicate FIT, with numbers of TPs and TNs 1004 and 800 (pan-cancer) and 93 and 800 (pancreatic). Critical values at 99% significance level, Bonferroni corrected for simultaneous testing of 53352 hypotheses. (b) Upper 99.9^th^ percentile of Subfigure a, with MC-simulation and exact computation bridged by geometric interpolations. For pancreatic cohort, an alternative interpolation with monotonically increasing geometric coefficient is included. (c) Ratio of subsequent differences for pan-cancer cohort, with a ratio inferred from the median of the S-shaped distribution function at AUC 0.88. (d) Ratio of subsequent differences for pancreatic cohort, with a non-constant geometric coefficient, connecting the exactly computed ratios with the inferred ratio through a monotonically increasing positive exponent function.

Determining the numbers *k*_0_, *k*_1_, *k*_2_, …, i.e. the numbers of distinct ROC-curves that yield an AUC-value, see Figure 1 for an illustrative example, can be achieved through the following recursion, which is stylized in pseudo-code in Algorithm 1. The recursive algorithm employs the recursive equality that is described in the following. Denote by (*u*·*v*) the number of permutations in which *u* rectangles, each with area 1/*nm* cf. Figure 1, can be subtracted under a maximum of *v* rectangle side lengths. For instance, the side length may be taken along the vertical axis, cf. Figure 1. The number of permutations (*u*·*v*) can be decomposed into two parts as per the equality

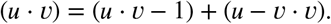

The first part of the decomposition are the permutations that use only *v*−1 side lengths, and the second part are the permutations that use all *v* side lengths which therefore equals the number of permutations (*u* − *v* · *v*).

The recursion terminates under the conditions (*u* · *v*) = 0 for all *u* < 0, i.e. there are nil permutations in which a negative number of rectangles can be subtracted, (*u* · 1) = 1, i.e. there is one permutation in which any number of rectangles can be subtracted under a maximum of only one side length, and (0 · *v*) = 1, i.e. there is one permutation in which nil rectangles can be subtracted under any maximum of rectangle side lengths. Further, a condition on the maximum of the number of side lengths along the axis perpendicular relative to the axis thus discussed is implemented, denoted by max_dim2 in the pseudo-code. While the authors view it as possible that a closed form expression for the numbers *k*_*i*_, *i* = 0, 1, …, *nm*, exists, we have not at time of writing been able to derive such; hence the recursive algorithm.

#### Algorithm 1

Recursive algorithm returning the number of ROC curves that yield an input AUC-value.

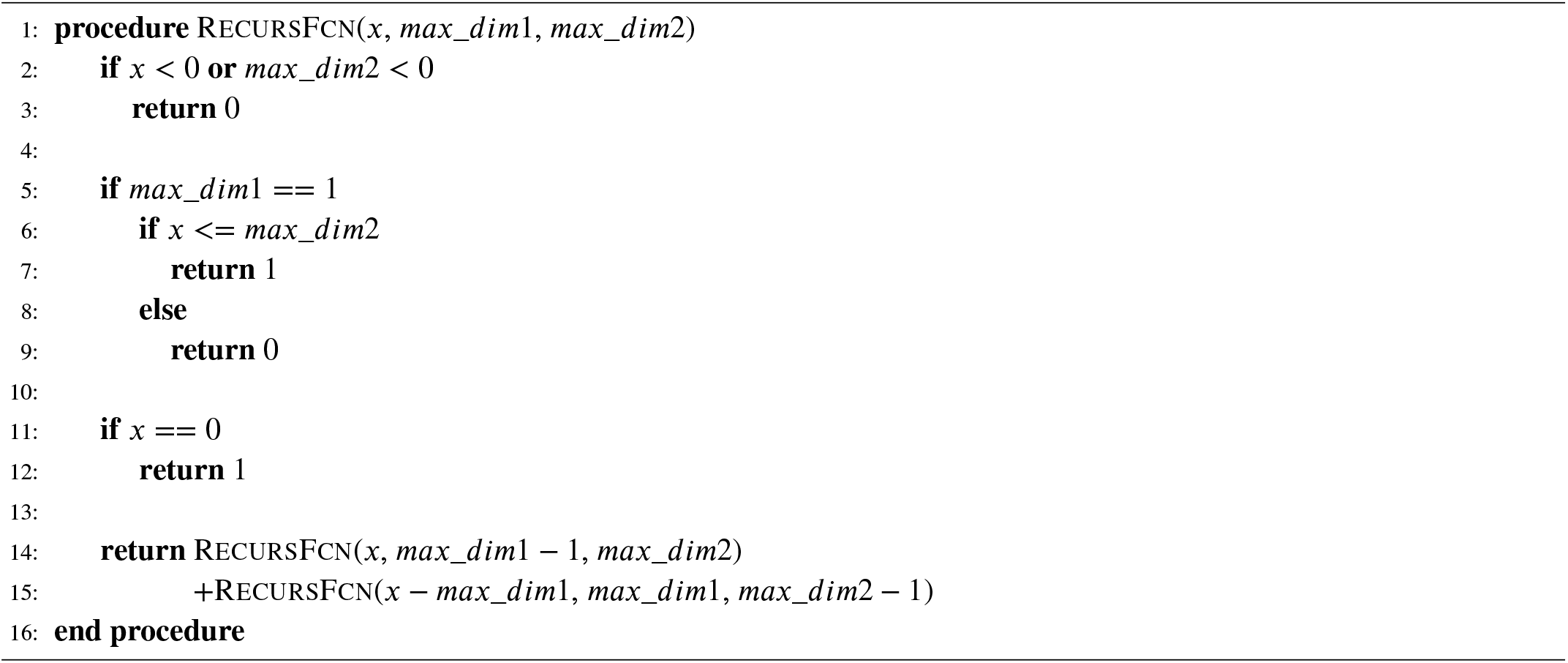

## 3 STATISTICAL HYPOTHESIS TESTING OF AUC-VALUES

Because the AUC-value is a number in the unit interval, a one-sided acceptance region with type-one error probability *α* is constructed through the interval [0, *c*] where *c* is the critical value satisfying *P* (*X* ≥ *c*) = *α* and *X* is the random test variable under the null-hypothesis^21^. In applications such as biomarker discovery, it is common to simultaneously consider numerous binary predictors. In particular, systematically forming composite biomarkers from constituent measurands often causes a combinatory proliferation, and thus AUC-values for millions of composite biomarkers may be simultaneously considered. As a result of correction for testing of multiple hypotheses through Bonferroni-correction or other method, the type-one error probability, *α*, is commonly tiny. For a numerical example, a statistical significance level of 99% and 10^8^ composite biomarkers yields a Bonferroni-corrected *α*-value of 10^−10^.

With AUC-values, estimation of the critical value through outright Monte Carlo-simulation will be especially taxing because each AUC-value requires simulation of observed values of TPs and TNs. Within the setting of a clinical trial the numbers of TPs and TNs could be between 100 and 1000 each, and to obtain an estimate of a critical value at *α* equalling 10^−10^, at least 10^10^ + 1 AUC-value simulations are needed. Hence, to obtain the most coarse estimate of the critical value through Monte Carlo-simulation, several trillion random numbers are often needed, and to obtain a more precise estimate perhaps a quadrillion random numbers would be desired in the presently discussed numerical example.

Furthermore, when the purpose of the Monte Carlo-simulation is estimation of statistical power or experimental design optimization, then simulations will typically need to be performed under a range of design parameter values, which will greatly compound the difficulties. For a numerical indication, at time of writing the authors are able to simulate 10^10^ AUC-values in 2.63 · 10^6^ seconds, or 30.4 days, at 400 cases and 400 controls using highly optimized algorithms in R (version 3.3.2)^22^. Simulation across a range of sample sizes and possibly other design parameters would multiply the required time several fold. Hence, statistical hypothesis testing of AUC-values is simple in principle, but the determination of the probability distribution tails is challenging computationally.

## 4 EXAMPLES

### 4.1 Illustration through simulated data

For illustrative purposes, a few numerical examples are provided. In these examples the observed values of TPs and TNs are each normally distributed with unit variance and some difference in their mean values. When the difference in means is taken to be equal to one, then the resulting AUC-value is on average about 0.76 with a standard deviation that depends largely on the number of TPs and TNs.

Figures 2a and 2b show probability distribution functions of the AUC-value when the differences in the means of the observed values of TPs and TNs are equal to one, and the number of TPs and TNs are *n* = *m* = 10 and *n* = *m* = 50 respectively. When the numbers of TNs and TPs are 10, then the AUC-value can attain 101 distinct values and consequently it is feasible to compute the probability of each AUC-value exactly, as per Section 2.1. Also, estimates of the probabilities obtained through Monte Carlo-simulation, as per Section 2.2, are shown. Alignment between the two are visually evident.

In Figure 2b, the numbers of TNs and TPs are 50, and consequently the AUC-value can attain 2501 distinct values, and with the computers used at time of writing computation of exact probabilities of all AUC-values, as per Section 2.1, was deemed infeasible. The 57 AUC-value probabilities shown in Figure 2b have a sum of about 5.05 · 10^−10^, and the turquoise line shows geometric interpolation, as per Section 2.3, between the left-most AUC-value probability and the 99^th^ percentile estimated through Monte Carlo-simulation. As examples, AUC-values 0.90, 0.95 and 0.99 have p-values 4.4 · 10^−4^, 4.3 · 10^−7^ and 8.5 · 10^−16^ respectively. Similarly, under Bonferroni-correction for 10^8^ hypotheses the critical value for a one-sided test at the 99% significance level is 0.9816, i.e. if any of the observed AUC-values is greater than 0.9816 then the null-hypothesis is rejected; it seems improbable that the observed value is an observation from the hypothesized probability distribution.

Figures 2c and 2d illustrate the degree of stability of the ratios of subsequent probability differences normalized by the corresponding ratios of the differences of numbers of ROC-curves per AUC-values, i.e. the normalized ratios (Δ*x*_*i*_/Δ*x*_*i*−1_)/(Δ*k*_*i*_/Δ*k*_*i*−1_), for the first 56 differences, which were the greatest number deemed feasible to compute when preparing the present example. As discussed in Section 2.3, when the TP and TN-distribution difference in means is zero the ratio is identically equal to unity. The constant *C*, discussed in Section 2.3, is estimated using the bisection method, and is plotted in the figures in dashed lines. Alignments between the stabilizing ratios and the constants *C* estimated using the bisection method are visually evident.

Because the distribution function curve, cf. Figure 2b, possesses an S-shape, i.e. first accelerates then decelerates, the ratio of subsequent probability differences is likely approximately constant only within an interval of limited length. However, as discussed in Section 2.3, the need to bridge exact computations and Monte Carlo-estimates is typically confined to the top or bottom percentile of the probability distribution, and within that finite region the approximation tends to be quite valid as illustrated in Figures 2c and 2d.

### 4.2 Illustration through biomarker data

Statistical hypothesis testing and determination of p-values of AUC-values under correction for multiple hypothesis testing is illustrated using the proteomic data set of^20^. The data encompass 39 proteins measured across 1004 individuals newly diagnosed with cancer and 812 healthy controls. The cancer diagnoses are breast, colorectal, esophagus, liver, lung, ovary, pancreas, and stomach cancer. In addition, a ninth pan-cancer diagnosis is formed by merging the eight cancer diagnoses. It may be noted that the aforementioned data source did not include statistical hypothesis tests or p-values. For ovarian cancer, we chose to only include female healthy controls.

Composite biomarkers are formed as per^23^. From the 39 proteins, 741 composite biomarkers are formed by combining the proteins into pairs. The 741 are further multiplied by 8 as a result of allowing positive classification when one or both of the proteins are down-regulated and when either or both of the proteins are up or down-regulated. By testing for the 9 diagnoses, a total of (39 choose 2) · 8 · 9 = 53352 composite biomarkers are simultaneously statistically hypothesis tested. P-values are adjusted as per the Bonferroni method. The null-hypothesis is that the biomarkers are on par with the colorectal cancer test FIT, which has AUC-value 0.88^24, p. 31^. As a commonly used screening test, the stool-based test FIT is a direct predicate IVD device for the colorectal cancer cohort, and also a relevant benchmark for the other cohorts that lack regulatory approved screening predicate IVD devices^25^. Rejection of the null-hypothesis is interpreted as evidence that, sources of error notwithstanding, the biomarker has AUC-value that is superior vis-à-vis the AUC-value of FIT.

For each of the 9 diagnoses, 58 AUC-values were computed exactly, i.e. 1, 1 − 1/*nm*, 1 − 2/*nm*, …, 1 − 57/*nm* where *n* and *m* are the numbers of TPs and TNs respectively. The 9 probability distributions were then interpolated geometrically, as per Section 2.3, to the 99.99^th^ percentiles which were determined by Monte Carlo-simulation of 10^8^ AUC-values per diagnosis. The distinctly large Monte Carlo-simulation was used because some of the cohorts have highly unbalanced numbers of TPs to TNs, which affects the geometric interpolation as discussed in Section 2.3 and illustrated in Figure 3.

In total, 26 cancer biomarkers were statistically significant at the 99%-level, meaning that they have an AUC-value that is higher than the highest AUC-value out of 53352 biomarkers on par with FIT would reasonably produce. The interpretation is that those biomarkers are, sources of error notwithstanding, superior relative to FIT. In this instance, separation of training and validation data does not constitute an issue because every biomarker is tested statistically; i.e. no training is conducted.

Table 1 shows Bonferroni corrected critical values relative to the null-hypothesis that the 53352 biomarkers have AUC-value on par with FIT. It is evident that tumor type cohorts that have larger sample sizes exhibit critical values that are lower than for cohorts that have relatively smaller sample sizes. Of the biomarkers tested, 26 are significantly better, at the 99% level, than FIT, including 4 liver, 14 ovarian, and 8 pancreatic cancer biomarkers. Table 2 details those 26 biomarkers; which proteins they use, whether the proteins are up or down regulated, whether up or down regulation of either protein or both is necessary for positive classification. A selection of significant biomarkers are shown graphically in Figure 4.

**Table 1.** Critical values for two-protein biomarkers. One-sided 99% critical values relative to the null-hypothesis that the biomarker has AUC-value on par with the colorectal cancer test FIT, which has AUC 0.88, Bonferroni-corrected for simultaneous testing of 53352 hypotheses. The numbers of TPs and TNs account for some missing values.

| Tumor type cohort | Number of<br>TPs TNs |  | Critical value<br>99%, correct'd | Number of<br>sign. biom. |
| --- | --- | --- | --- | --- |
| Breast | 209 | 800 | 0.942 | 0 |
| Colorectal | 388 | 800 | 0.930 | 0 |
| Esophagus | 45 | 800 | 0.978 | 0 |
| Liver | 44 | 800 | 0.978 | 4 |
| Lung | 104 | 800 | 0.959 | 0 |
| Ovary | 54 | 372 | 0.974 | 14 |
| Pancreas | 93 | 800 | 0.961 | 8 |
| Stomach | 68 | 800 | 0.969 | 0 |
| Pan-cancer | 1004 | 800 | 0.919 | 0 |

**Table 2.** Two-protein blood biomarkers for liver, ovarian and pancreatic cancers. Details of the 26 cancer biomarkers that have AUC values greater than the critical values in Table 1. Proteomic data set from^20^ and composite biomarkers are formed as per^23^. Up arrow, ↑, the protein is up-regulated among donors who have cancer relative to the healthy controls. Down arrow, ↓, the protein is down-regulated among cancers relative to controls. ‡ Whether it is necessary for positive classification that either or both of the proteins are up or down-regulated. § The pAUC-value is computed in the interval [0, 0.2]. ¶ The p-value is Bonferroni corrected for simultaneous hypothesis testing of 53352 biomarkers, under the null-hypothesis that each are on par with the colorectal cancer test FIT which has AUC 0.88^24, p. 21^.

| Tumor type | Proteins* | Regulation‡ | AUC | pAUC§ | P-value¶ |
| --- | --- | --- | --- | --- | --- |
| Liver | HGF↑ OPN↑ | Either | 0.983 | 0.183 | $4.33 \cdot 10^{-4}$ |
| | AFP↑ OPN↑ | Either | 0.982 | 0.187 | $7.85 \cdot 10^{-4}$ |
| | HGF↑ PRL↑ | Either | 0.979 | 0.182 | $7.03 \cdot 10^{-3}$ |
| | GDF15↑ IL-8↑ | Both | 0.979 | 0.179 | $8.73 \cdot 10^{-3}$ |
| Ovary | CA-125↑ PRL↑ | Either | 0.987 | 0.194 | $1.63 \cdot 10^{-6}$ |
| | CA-125↑ TIMP-1↑ | Either | 0.984 | 0.186 | $6.67 \cdot 10^{-6}$ |
| | CA-125↑ TSP2↓ | Either | 0.985 | 0.192 | $2.32 \cdot 10^{-5}$ |
| | CA-125↑ IL-6↑ | Either | 0.982 | 0.184 | $8.12 \cdot 10^{-5}$ |
| | CA125↑ IL-6↑ | Both | 0.981 | 0.181 | $1.76 \cdot 10^{-4}$ |
| | PRL↑ TIMP-1↑ | Either | 0.980 | 0.180 | $3.66 \cdot 10^{-4}$ |
| | CA-125↑ GDF15↑ | Either | 0.978 | 0.182 | $1.22 \cdot 10^{-3}$ |
| | CA-125↑ CEA↓ | Either | 0.977 | 0.179 | $2.40 \cdot 10^{-3}$ |
| | CA-125↑ OPN↑ | Both | 0.977 | 0.182 | $2.61 \cdot 10^{-3}$ |
| | CA-125↑ TGF-α↑ | Either | 0.976 | 0.184 | $2.76 \cdot 10^{-3}$ |
| | CA-125↑ OPN↑ | Either | 0.976 | 0.182 | $3.16 \cdot 10^{-3}$ |
| | CA-125↑ sFas↓ | Either | 0.976 | 0.176 | $4.23 \cdot 10^{-3}$ |
| | IL-6↑ sFas↓ | Both | 0.975 | 0.178 | $6.06 \cdot 10^{-3}$ |
| | CA-125↑ ENG↑ | Either | 0.974 | 0.177 | $9.03 \cdot 10^{-3}$ |
| Pancreas | CA19-9↑ sErbB2↑ | Either | 0.971 | 0.176 | $1.88 \cdot 10^{-5}$ |
| | CA19-9↑ GDF15↑ | Either | 0.967 | 0.172 | $3.68 \cdot 10^{-4}$ |
| | CA19-9↑ OPN↑ | Either | 0.966 | 0.171 | $8.14 \cdot 10^{-4}$ |
| | CA19-9↑ IL-6↑ | Either | 0.966 | 0.172 | $1.05 \cdot 10^{-3}$ |
| | CA19-9↑ TIMP-2↑ | Either | 0.964 | 0.172 | $2.22 \cdot 10^{-3}$ |
| | CA19-9↑ IL-8↑ | Either | 0.962 | 0.168 | $8.33 \cdot 10^{-3}$ |
| | CA19-9↑ PRL↑ | Either | 0.962 | 0.170 | $8.89 \cdot 10^{-3}$ |
| | CA19-9↑ HGF↑ | Either | 0.961 | 0.173 | $9.81 \cdot 10^{-3}$ |

**Figure 4.**
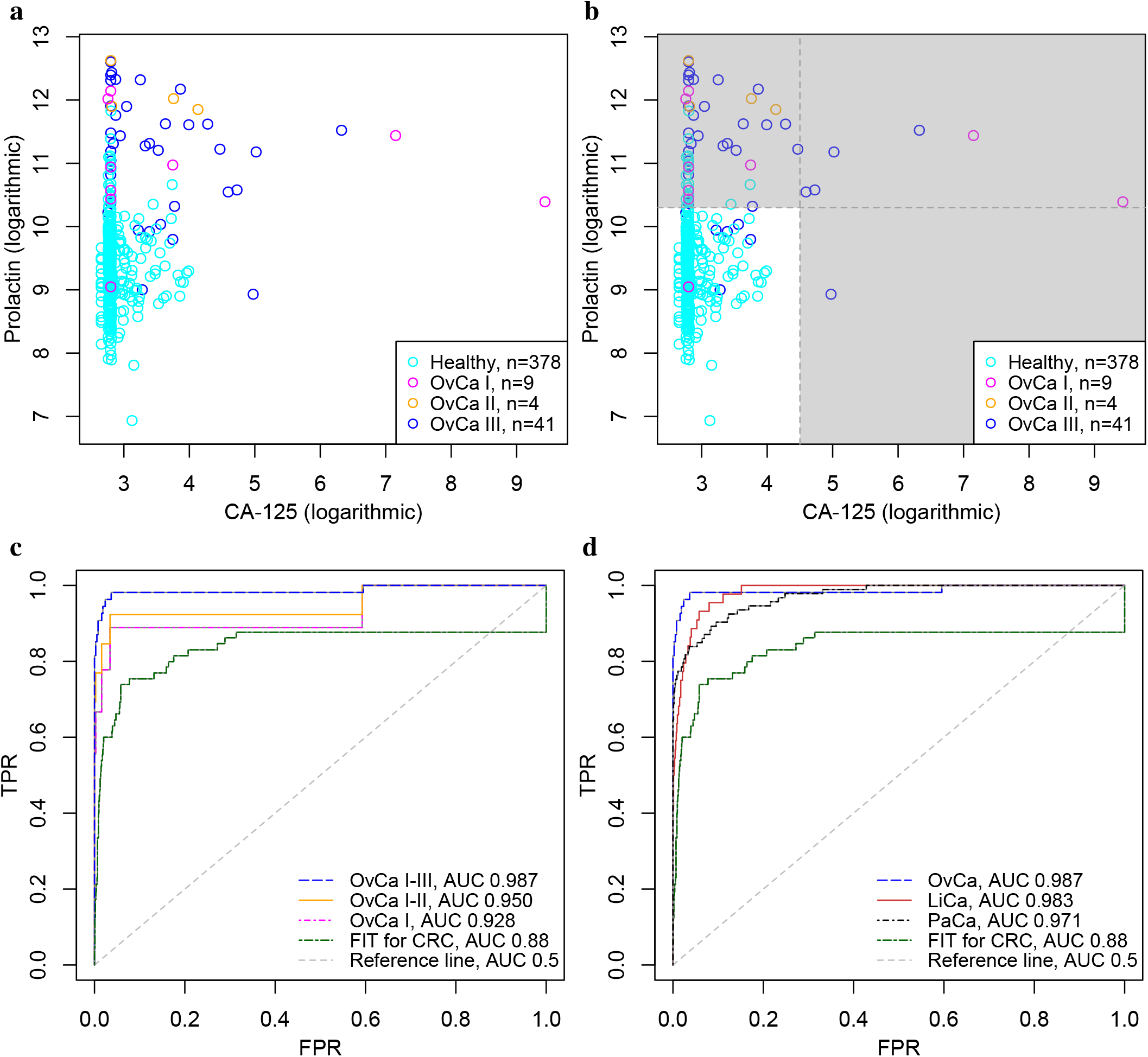
Biomarkers for detection of ovarian, liver and pancreatic cancers, that are superior vis-Á-vis benchmark predicate device FIT at the 99% significance level. (a) Scatter plot of CancerSeek data, CA-125 and Prolactin, per OvCa stage (I, II and III) and female healthy controls. (b) Scatter plot similar to Subfigure a, with cut-offs at 4.5 and 10.3; illustrating positive and negative classification regions. (c) ROC-curves for CA-125 and Prolactin as a biomarker for detection of OvCa, per stages. Stronger biomarker signals for more advanced cancers are evident. For reference, FIT for CRC and the diagonal line is included. (d) ROC-curves for biomarkers for detection of OvCa (CA-125, Prolactin), LiCa (HGF, Osteopontin) and PaCa (CA 19-9, sErbB2). FIT and the diagonal line included for reference.

## 5 DISCUSSION

Historically, AUC-value probability distributions have been approximated under assumptions of asymptotic normality, allowing t-tests or similar^16,17^; however the approach provides inadequate accuracy in the tails of the probability distribution where critical values under correction for multiple hypotheses are situated. With exact computation of AUC-value probabilities, impeccable precision is obtainable; permitting accurate statistical inference. In biomarker development, if critical values are too low the type-one error probability will be too high; effectively yielding false expectations. If critical values are too high, then the type-two error probability will be too high; effectively barring identification of potentially valid biomarkers. Using the exact AUC-value probability distribution, the aforementioned risks are avoided, all while allowing for computation of exact p-values.

In order to provide reasonable assurance that a study has sufficiently large sample sizes to demonstrate a putative effect, experimental design optimization is warranted. The computational challenges inherent to Monte Carlo-simulation are greatly compounded when critical values under a multidimensional set of parameter values need to be estimated. With the proposed method of exact computation of AUC-value probabilities paired with Monte Carlo-simulations and geometric interpolation, a method is obtained that is both computationally feasible and relatively precise.

## ^0^Abbreviations

UC: area under curve
*iid*: independent and identically distributed
IVD: *in vitro* diagnostic
ROC: receiver operating characteristic
TN: true negative
TP: true positive

## DATA AVAILABILITY STATEMENT

Code for simulation of AUC-value data available on GitHub https://github.com/TSResearchGroupUU/, Biomarker data available online via^20^.

